# Unclearing Microscopy

**DOI:** 10.1101/2022.11.29.518361

**Authors:** Ons M’Saad, Michael Shribak, Joerg Bewersdorf

**Author notes:** Corresponding author. (J.B.).

## Abstract

The spatial resolution and contrast sensitivity of the human eye is limited, restricting our ability to directly see subcellular structures. We report a new principle for unaided eye cellular visualization in a method we call Unclearing Microscopy. By expanding cells and tissue >8,000 volumetrically and opaquing their bulk with light-scattering molecules of sufficient density, cell microstructure can now be discerned with a contrast visible to the unaided eye. We further inspect uncleared samples with transmitted light microscopy modalities and prove that 3D ultrastructural features, previously accessible only with super-resolution fluorescence or electron microscopy methods, can now be visualized with simple magnification optics alone.

## Main

Fluorescence microscopy is one of the major light microscopy modalities in the life sciences. It uses refractive lenses to magnify objects of interest and relies on fluorescent labels that induce a highly sensitive contrast (Ellinger, 1940). Fluorescence can be detected in single-molecule amounts, making it an ideal contrast mechanism to image the distribution of proteins on the nanoscale (Shashkova and Leake, 2017). In super-resolution microscopy (SRM), selectively detecting subsets of fluorophores in a diffraction-limited volume allows up to single-digit isotropic spatial resolution in cells and tissue, revealing precise subcellular protein distributions on the nanoscale (Baddeley and Bewersdorf, 2018; Gwosch et al., 2020; Liu et al., 2022).

The physical magnification of a fixed biological sample itself is also possible in a technique known as Expansion Microscopy (ExM) (Chen et al., 2015). Here a sample is labeled with fluorescent molecules, crosslinked to a polyanionic hydrogel, mechanically homogenized, and swelled 4-fold or more linearly in low ionic strength solutions. To capture nanoscale information, expanded samples are usually imaged on a fluorescence microscope and the total image magnification is the product between both physical and optical magnifications (Wassie et al., 2019; Gallagher and Zhao, 2021; Wang et al., 2022). Using high-end confocal imaging systems, the apparent lateral spatial resolution achieved with ExM techniques usually ranges from ~15 nm to ~70 nm, depending on expansion factor, sampling, and labeling density.

In our previous work, we demonstrated that by expanding biological specimens ~20-fold linearly while simultaneously retaining the protein content, bulk- (pan-) staining of reactive amino acids resolves local protein densities and reveals ultrastructural context by standard confocal fluorescence microscopy (M’Saad and Bewersdorf, 2020; M’Saad et al. 2022). We call this method pan-Expansion Microscopy (pan-ExM). Analogous to electron microscopy (EM), subcellular compartments can now be imaged in their entirety, without the need for specific labels and with a lateral spatial resolution of ~15 nm. Our pan-staining concept has been since adapted by many labs to examine contextual spatial information in a multitude of biological systems, and at different expansion levels, demonstrating the wide utility of this contrasting approach (Nijenhuis et al., 2021; Sim et al., 2021; Klimas et al., 2021; Damstra et al., 2022; Laporte et al., 2022; Shaib et al., 2022; Hinterndorfer et al., 2022; Rashpa and Brochet, 2022; Pacheco et al., 2021; Suen et al. 2022).

While fluorescence microscopy was our default approach to visualize biological material at high spatial resolution, we asked whether cells (~50 μm) and their microstructures (~5 μm) can be discerned with the human eye alone without relying on additional optical manipulation. Such development would represent a new way of studying biological microstructures, circumventing the need for an intermediary apparatus.

In fact, if a typical HeLa cell of typical ~40 μm width is expanded 20-fold linearly, its width grows to 0.8 mm. Since the human eye has a resolution of 35-60 arcseconds (Anderson et al., 1991), it could in principle, at a distance of ~30 cm, resolve objects at the ~50 μm scale, corresponding to fine microstructures of a 20-fold linearly expanded HeLa cell. However, accompanying this expansion, the protein content of a single cell (~1.1 mM) (Milo and Phillips, 2015) will be diluted 20^3^=8,000-fold to ~140 nM. This poses a severe hurdle since the limited contrast sensitivity of the human eye (Kimpe & Tuytschaever, 2006) requires even under ideal conditions at least a 410-fold higher concentration (58 μM) of a highly absorptive dye (molar absorption coefficient of 10^5^ M^-1^ cm^-1^) to detect a 50-μm wide and thick object (see **Supplementary Information**; **Fig. 1a**).

**Figure 1:**
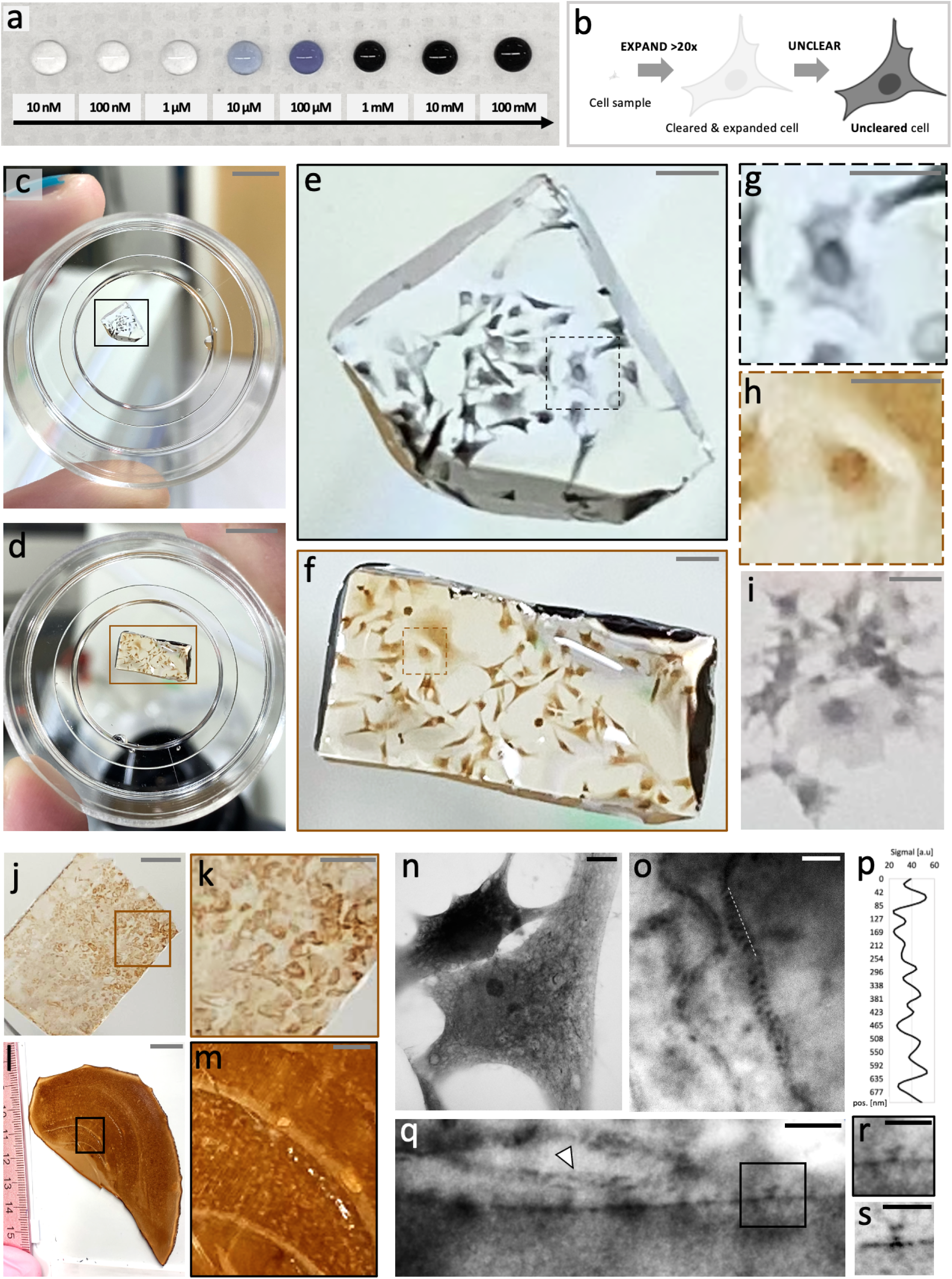
Unclearing by chromogen deposition reveals cellular microstructure to the unaided eye. **a**, Evans blue dye at indicated concentrations. **b**, Schematic of the Unclearing concept. **c**, **d**, Metallic silver (**c**) and DAB (**d**) uncleared HeLa cells on a glass bottom dish, imaged with a cell phone camera. **e**, **f**, Magnified views of the areas outlined by the black and brown boxes in **c** and **d**, respectively. **g**, **h**, Magnified views of the areas outlined by the dashed black and brown boxes in **e** and **f**, respectively. **i**, Metallic silver uncleared COS-7 cells imaged with a cell phone camera. **j**, DAB uncleared kidney tissue imaged with a cell phone camera. **k**, Magnified view of the area outlined by the brown box in **j** showing kidney tubules. **l**, DAB uncleared 5xFAD mouse hippocampus imaged with a cell phone camera. **m**, Magnified view of the area outlined by the black box in **l**. **n**, DAB uncleared HeLa cells imaged with transmitted light microscopy. **o**, DAB uncleared mitochondria in a HeLa cell showing visible cristae. **p**, Line profile along the dashed line shown in **o**. **q**, Zoomed in image of a nucleus border in a DAB uncleared HeLa cell showing mitochondria (white arrow) and a nuclear pore complex (NPC) (black box). **r**, Magnified view of the area outlined by the black box in **q**. **s**, Fluorescently pan-stained NPC which is qualitatively similar to the uncleared NPC shown in **r**. Scale bars shown in gray are not corrected for the expansion factor. Black scale bars are corrected for the determined linear expansion factor of 14.0. Scale bars, (**c**, **d**, **j**) 5 mm, (**e**, **f**) 1 mm, (**g**, **h, i**) 0.5 mm, (**k**, **m**) 2 mm, (**l**) 10 mm, (**n**) 5 μm, (**o**, **q**) 500 nm, (**r**) 250 nm,

We therefore reasoned that combining physical magnification of cells with amplification of the pan stain could result in a visible cellular contrast, with microstructures ultimately resolvable by the unaided eye. We speculated that if we pan-stained expanded cells with peroxidases or photoinitiators and amplified the underlying chromogenic substrate signal 10^3^-10^6^-fold, the resulting ~0.14-140 mM pigment concentration would be easily detected by the human eye (**Fig. 1a**). Earlier studies determined that horseradish peroxidase (HRP) amplification coupled with 3,3’-diaminobenzidine (DAB) deposition (Adams, 1991) and photopolymerization-based signal amplification (PBA) (Hansen et al., 2008) can yield amplification degrees of equivalent orders of magnitude, strengthening the feasibility of our approach.

Here, we introduce Unclearing Microscopy, a method where biological samples are expanded, pan-stained, and their pan stain amplified (uncleared) to yield a chromogenic product visible by the unaided eye. We further show imaging of uncleared cells and tissue samples with transmitted light microscopy, demonstrating that ultrastructural cell and tissue features such as mitochondria cristae and neuronal synapses can be revealed without complex super-resolution instrumentation or electron microscopy, using magnification optics alone.

## Results

### Unclearing by chromogen deposition

**Figure 1b** outlines the basic workflow of Unclearing Microscopy. Cells are first expanded ~20-fold with pan-ExM, pan-stained with amine-reactive NHS-ester ligands conjugated to biotin (NHS-PEG4-Biotin), and incubated with streptavidin conjugated to horseradish peroxidase (ST-HRP). The hydrogel sample is then sectioned to a thickness of ~1 mm and uncleared with chromogenic substrates 3,3’-diaminobenzidine (DAB) or metallic silver. The sample is finally washed with water and visualized directly.

**Figure 1c–i** confirms the validity of our concept. Taken with a standard cell phone camera under typical room light conditions, uncleared gels reveal individual mammalian cells and their microstructures. We found that metallic silver and DAB substrates yielded strong black and brown chromogenic contrast, respectively. We can easily discern cellular boundaries, cell contact sites, the cell cytosol, and the nucleus by eye. Similarly, when applying the same principle to 5xFAD mouse brain and mouse kidney tissue sections (**Supplementary Fig. 1**), unclearing reveals individual tubules in the renal cortex (**Fig. 1j–k**) and the different layers of the hippocampus (**Fig. 1l–m**) by the unaided eye.

Imaging our uncleared samples with conventional phase contrast microscopy, revealed ultrastructural features at higher magnification and contrast that in the past could only be seen by super-resolution fluorescence or electron microscopy. **Fig. 1n** shows whole HeLa cells uncleared with DAB. Mitochondria cristae, the nuclear envelope, and nuclear pore complexes (NPCs) can also be discerned (**Fig. 1o–s**). These images resemble those acquired with fluorescently pan-stained cells in pan-ExM but have the benefits of (1) no reliance on photon emitters which are susceptible to bleaching and (2) minimal degradation of staining over time. The fact that we can resolve mitochondria cristae (**Fig. 1o–p**) which are only ~70 nm apart in unexpanded cells (Stephan et al., 2019), suggests that blurring of features from signal amplification is minimal on that scale.

### Unclearing by photopolymerization

Next, we investigated whether amplifying the pan-stain using *in situ* polymerization of a highly crosslinked photopolymer could also render cells visible by eye and perhaps even confer a certain structural sturdiness to the enlarged cells in question. Such a technique would represent an orthogonal method of tissue unclearing, widening its applicability in biological studies.

To unclear biological samples using *in situ* polymerization, we used a strategy called photopolymerization-based signal amplification (PBA). PBA is commonly used to amplify signals from antibody-antigen interactions in immunoassays (Hoang et al., 2021). In these assays, antigens bind to immobilized antibodies and are then mixed with photoinitiator-conjugated capture antibodies. These also bind the antigens and when irradiated with light, in the presence of hydrogel monomers and catalysts, initiate free-radical polymerization. PBA is thereby spatially confined to the photoinitiator location and yields a polymer film of ~10-250 nm thickness, depending on hydrogel monomer composition and photoinitiator density (Hansen et al., 2008).

Most PBA methods use eosin as a photoinitiator because it is a high triplet yield fluorophore that can partially regenerate in oxygen (Avens and Bowman, 2009). Coupled with tertiary amine methyl diethanolamine (MDEA) and under 530-nm green light irradiation, eosin and MDEA undergo energy transfer and initiate free-radical polymerization in the presence of common hydrogel monomers and crosslinkers (**Fig. 2a**) (Hansen et al., 2008; Avens and Bowman, 2009).

**Figure 2:**
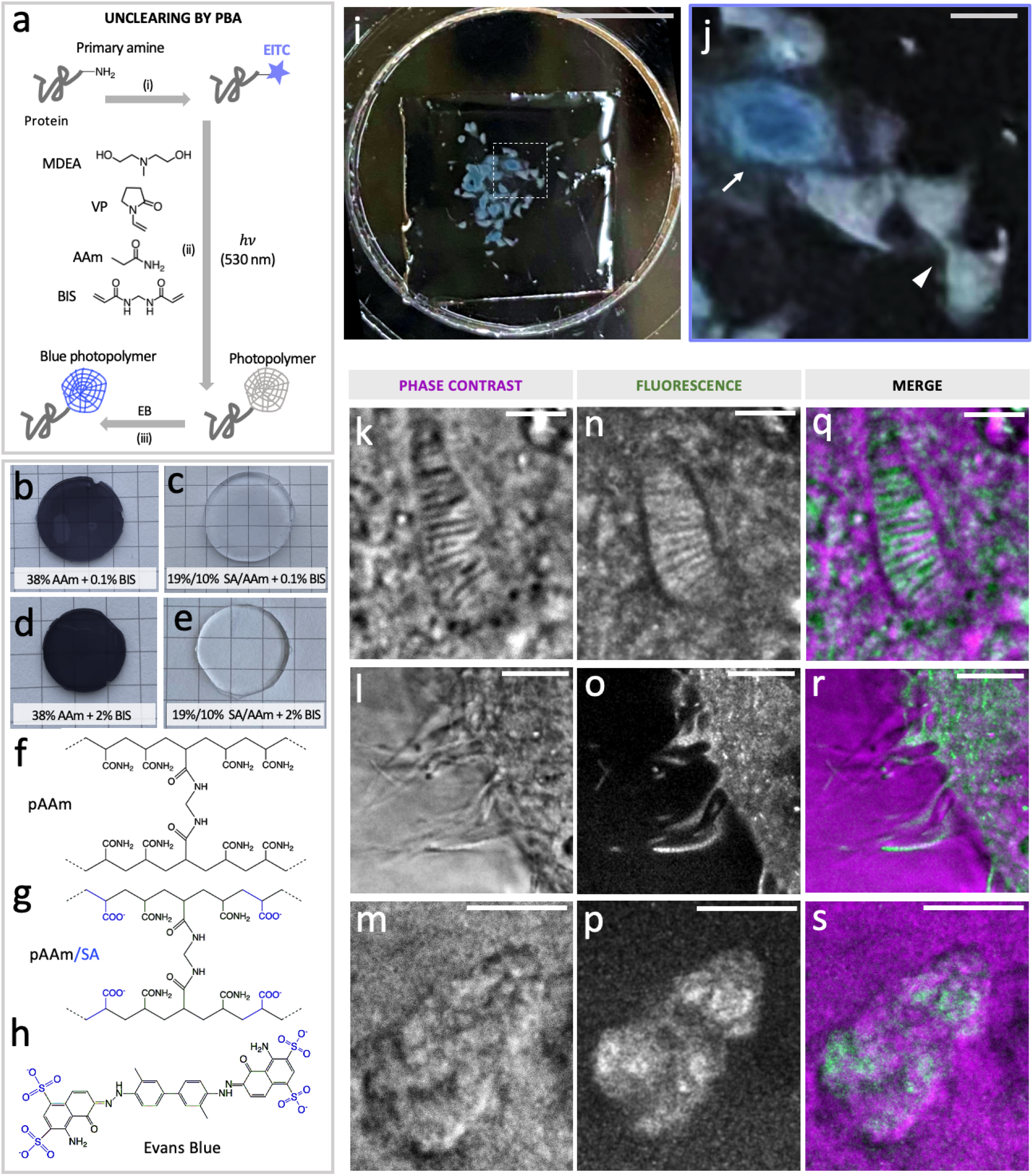
Unclearing by in situ photopolymerization. **a,** The workflow on unclearing using photopolymerization-based signal amplification (PBA): proteins in a pan-ExM processed sample are (i) pan-stained by eosin 5-isothiocynate (EITC), (ii) incubated in a solution of acrylamide monomers (AAm) and N-vinylpyrrolidone (VP), crosslinker N,N’-methylenebisacrylamide (BIS), and tertiary amine methyl diethanolamine (MDEA), (iii) photopolymerized by exposure to 530-nm LED light, and (iv) stained with Evans Blue (EB). **b-e**, Evans Blue, an anionic dye, efficiently stains neutral polyacrylamide hydrogels (**b** and **d**) but not anionic poly(acrylamide/sodium acrylate) co-polymers (**c** and **e**), regardless of hydrogel crosslinker concentration. **f**, Chemical structure of polyacrylamide (pAAm) polymer. **g**, Chemical structure of poly(acrylamide/sodium acrylate) co-polymer (pAAm/SA) (blue: negatively charged carboxylic groups). **h**, Chemical structure of Evans Blue dye (blue: negatively charged sulfate groups that repel carboxylic groups in anionic pAAm/SA hydrogels and prevent staining of the hydrogel). **i**, PBA uncleared HeLa cells stained with Evans Blue and imaged with a cell phone camera. **j**, Magnified view of the area outlined by the dashed white box in **i** showing cells in interphase (arrow) and cytokinesis (arrowhead). **k-m**, Phase contrast images of a mitochondrion (**k**), filipodia (**l**), and a nucleolus (**m**) in PBA uncleared HeLa cells. **n-p**, Same areas as in **k-m**, showing fluorescent NHS ester pan-staining. **q-s**, Overlays of **k-m** and **n-p**. Scale bars in gray are not corrected for the expansion factor. White scale bars are corrected for the determined linear expansion factors of 17.5-20.4. Scale bars, (**i**) 5 mm, (**j**), 0.5 mm, (**k, n, q**), 250 nm, (**l, o, r**, **m, p, s**), 1 μm.

PBA has also been demonstrated to amplify fluorescence signals in several immunofluorescence assays (Avens et al., 2011a; 2011b). For example, one study used eosin light-initiated polyethylene (glycol) diacrylate (PEGDA) polymer films to entrap fluorescent nanoparticles onto microtubules, resulting in up to ~10^3^-fold amplification of fluorescence signal (Avens et al., 2011b).

Here, we investigated whether expanded cells pan-stained with eosin can be used to synthesize a densely crosslinked polyacrylamide hydrogel *in situ*. The photopolymer would be ideally capable of entrapping small chromogenic dyes, amplifying the pan-stain accordingly.

Since the cell protein content is significantly diluted following sample expansion (~140 nM), we first tested if eosin surface densities of equivalent concentrations can still initiate photopolymerization. In a solution of 38% (w/v) acrylamide (AAm), 35 mM N-vinylpyrrolidone (VP), 2% (w/w) methylene bisacrylamide (BIS), and 210 mM MDEA, we added eosin in concentrations of 0.03 μM to 12 μM (**Supplementary Fig. 2**). The aforementioned eosin concentrations correspond to that of proteins in a 20-fold expanded cell had they originally been separated by 5 nm to 40 nm respectively. Irradiating these solutions with 530-nm light with intensities between 12 and 55 mW/cm^2^, we confirmed that even at 0.03 μM eosin concentration, a photopolymer can still form, with increased thickness when the gelation solution is purged with nitrogen (**Supplementary Fig. 2f**). This finding is consistent with a previous report which determined that the eosin threshold concentration for PEGDA and acrylamide photopolymers is 0.03 μM for 5% conversion in 20 min (Avens et al., 2008).

Next, we investigated if the resulting photopolymer can be stained with chromogenic dyes. It has been previously shown that Evans Blue (EB) can efficiently stain PEGDA gels (Lilly et al., 2014). We questioned whether EB could also stain polyacrylamide (pAAm, **Fig. 2f**) and whether it does so without also staining the underlying expansion poly(acrylamide/sodium acrylate) hydrogel (pAAm/SA, **Fig. 2g**). We found that indeed EB stains pAAm gels and not pAAm/SA gels (**Fig. 2b–e**), even when the BIS crosslinker concentration is varied from 0.1% to 2% (w/w) to account for crosslinking density. We hypothesize that this is because EB is sulfonated (**Fig. 2h**) and thus repelled by carboxylic groups in pAAm/SA gels. This finding is also true for the sulfonated textile dye Direct Red 81 (**Supplementary Fig. 3**). However, we chose to use EB in our subsequent experimentation because of its superior visible contrast.

Ensuingly, we developed a workflow for unclearing biological samples by PBA. In brief, samples are expanded according to the pan-ExM protocol, pan-stained with amine-reactive eosin-5-isothiocyanate, sectioned to ~1 mm thickness, incubated in a hydrogel monomer solution containing MDEA and high concentration of BIS crosslinker, photo-irradiated with 530-nm light, washed with water, optionally stained with EB, and visualized directly.

Using this protocol, we observed uncleared cells with the unaided eye (**Fig. 2i–j**). Even without chromogenic EB staining, photopolymers are still visible against a dark background (**Supplementary Fig. 4**). This is likely caused by the significant polymer refractive index variations between the photopolymer and the expansion hydrogel (n~1.33). Like samples uncleared with chromogen deposition, we can distinguish between the cytosol and cell nucleus (**Fig. 2i–j**), delineate cell boundaries, and classify cells by their shapes and interactions.

Additionally, we can physically excise individual cells with tweezers and manipulate them mechanically. This is due to the mechanical sturdiness of the underlying photopolymer compared to the more delicate expansion hydrogel.

Moreover, PBA uncleared samples are compatible with phase contrast microscopy. We can observe the fine ultrastructures of mitochondria cristae (**Fig. 2k**), filopodia (**Fig. 2l**) as well as nucleoli (**Fig. 2m**). Co-staining with NHS ester fluorescent dyes and overlaying phase contrast with fluorescence confocal images shows that these structures fully overlap, with confocal fluorescence imaging offering better optical sectioning.

### OI-DIC reveals EM-like 3D ultrastructural features in uncleared brain tissue

PBA unclearing is also applicable to mouse brain tissue sections. With conventional phase contrast imaging, we were able to resolve the fine structure of cortical neuron bodies as well as putative axons (**Supplementary Fig. 5**). However, we were unable to resolve synaptic densities and distinguish between neighboring neurites as we have demonstrated with pan-ExM-t, a fluorescence ultrastructural imaging method (M’Saad et al. 2022).

To this end, we decided to image ~1 mm-thick expanded and uncleared brain tissue sections with an Orientation-Independent Differential Interference Contrast (OI-DIC) microscope, a quantitative phase contrast transmitted light imaging modality with 3D optical sectioning capabilities (Shribak and Inoué, 2006).

**Figure 3** demonstrates the compatibility of OI-DIC with PBA uncleared tissue. Hallmark ultrastructural features in cortical mouse brain tissue, including the fine structure of mitochondria (**Fig. 3a–b** and **Fig. 3u–v**), putative ER tubules (**Fig. 3a** and **Fig. 3s**), spine apparatuses (**Fig. 3a**), nucleoli (**Fig. 3r**), and putative synapses (**Fig. 3a–m** and **Fig. 3w–aa**) are now visible with transmitted light.

**Figure 3:**
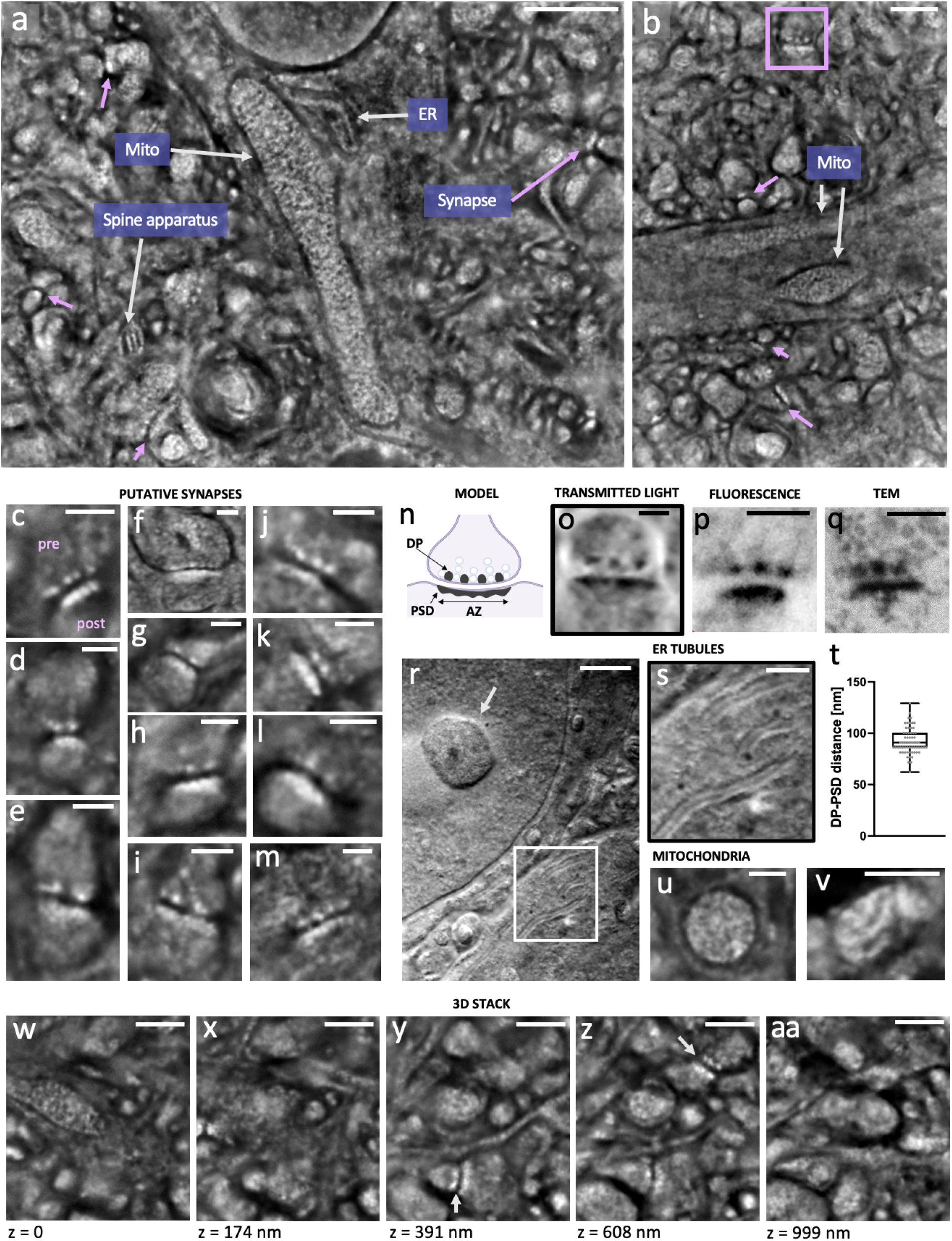
EM-like 3D ultrastructural features in uncleared brain tissue revealed by OI-DIC microscopy. **a**, **b**, Ultrastructural features such as mitochondria, putative endoplasmic reticulum (ER), spine apparatuses, putative synapses (lavender arrows and box) are visible in PBA uncleared mouse brain tissue when imaged with OI-DIC microscopy. **c-m**, Putative excitatory synapses determined by their characteristic synaptic densities: dense projects (DPs) in the presynaptic compartment (pre in **c**) and postsynaptic densities (PSDs) in the postsynaptic compartment (post in **c**). **n**, Schematic of a synapse highlighting synaptic densities. **o**, PBA uncleared putative synapse (inverted color table). **p**, fluorescence pan-ExM-t image of a putative synapse. **q**, Transmission Electron Microscopy (TEM) image of a PTA-stained putative synapse prepared as described in M’Saad et al., 2022. **r**, DIC image of a PBA uncleared brain tissue section showing a nucleolus (arrow) and putative ER tubules (box). **s**, Magnified image in the white box in **r. t**, DP-PSD distances (n = 63 measurements in 2 independent samples). **u**, **v**, Examples of PBA uncleared mitochondria in brain tissue. **w-aa**, z stack highlighting the 3D optical sectioning capabilities of OI-DIC. Arrows in **y** and **z** point to putative synapses. All OI-DIC images are Riesz-transformed. Scale bars are corrected for the determined linear expansion factor of 23.0. Scale bars, (**a**, **b**) 1 μm, (**c-m**) 200 nm, (**o-q**) 300 nm, (**r**) 250 nm, (**s**) 100 nm, (**u, v**) 250 nm, (**w-aa**) 500 nm.

Similar to pan-ExM-t, synaptic compartments can be identified by their characteristic synaptic densities (**Fig. 3c–m**), with dense projections (DPs) demarcating the presynaptic compartment and a macular postsynaptic density (PSD) representing the postsynaptic compartment. This characteristic pattern is almost identical to that visualized by confocal fluorescence microscopy of pan-ExM-t samples and transmission electron microscopy (TEM) of brain tissue stained with phosphotungstic acid (PTA) (**Fig. 3n–q**). Further, we found that the expansion-corrected DP-PSD distance in one experiment is 92.2 ± 12.7 nm (mean ± s.d, n = 63 measurements; **Fig. 3r**), consistent with that obtained in pan-ExM-t samples of the same experiment (81.6 ± 13.6 nm; mean ± s.d, n = 91 measurements) and in agreement with previous reports (Valtschanoff and Weinberg, 2001; Acuna et al., 2016; Hao et al., 2021).

Importantly, OI-DIC permits effective optical sectioning (**Fig. 3w–aa**) of PBA-uncleared specimen, paving the way to an era where neural circuit reconstruction with transmitted light comes within the realm of possibility.

## Discussion

It is indisputable that tissue *clearing* represents a revolution in the field of microscopy (Richardson and Lichtman, 2016). When biological tissue is cleared (i.e. transparentized and depigmented), limitations imposed by light scattering molecules (e.g, lipids, lipofuscin etc.) are removed, allowing for volumetric imaging of thick specimens on conventional light microscopes. While sample clearing has long been considered a desirable outcome of Expansion Microscopy (Chen et al., 2015) and is in fact an inherent property of this method, we show in this work that the opposite - *unclearing* (opaquing) - provides an equally exciting opportunity: unclearing structures of interest in a targeted manner generates contrast that reveals them in expanded biological samples without the need for complex instrumentation.

In fact, the data presented demonstrates that by sufficient sample enlargement and signal amplification, cellular microstructures can be revealed without the need for *any* optical instrumentation. While the demonstrated resolution is too small to reveal nanoscopic details with the unaided eye, larger linear expansion factors of 100-fold to 1000-fold in the future will render cellular organelles like mitochondria (~500 nm) large enough to see without supplementary optical magnification (e.g., ~100 μm after 200-fold linear expansion). While enticing, it is important to be cognizant of reaction kinetics when experimenting with larger expansion factors. For example, in the case of the SN2 nucleophilic substitution reaction of NHS ester with primary amines, the recommended minimum concentration of primary amines before NHS ester hydrolysis becomes dominant is 10 μM (Smith, 2006). Intermediate amplification steps with streptavidin and biotin (Chu et al., 2013) at expansion levels of 20-fold can help mediate this problem. In fact, the high affinity (KD ~10^-15^ M) of biotin for streptavidin indicates that the latter can be detected at femto- and picomolar concentrations (Delgadillo et al., 2019).

Moreover, our data demonstrates a conceptually new way of resolving ultrastructural details. Simple, laser-free optical systems such as widely available phase contrast microscopes or more advanced OI-DIC instruments can now reveal structures at the nanoscale in cells and tissues. We have shown here transmitted light images of nucleoli, mitochondria, filopodia, NPCs, ER tubules, synaptic densities, and spine apparatuses at sub-70 nm resolution. We have demonstrated unclearing by two methods: deposition of chromogens and targeted photopolymerization. However, other amplification strategies such as nucleic acid-based ones (Saka et al., 2019) should be equally feasible. We have shown ultrastructural cell and tissue imaging by amplifying the pan-stain, but our approach can also be adapted to imaging specific molecules using antibody labels conjugated to HRP or eosin. To extend Unclearing Microscopy to multiplexing in the future, we envision using differently colored chromogens (e.g., red and blue), different catalytic agents (e.g., alkaline phosphatase (AP) (Stack et al., 2014)), and different water-soluble photoinitiators (e.g., phenyl-2,4,6-trimethylbenzoylphosphinate (LAP) which is activated with UV light (Tomal and Ortyl, 2020)).

In contrast to established sub-diffraction optical microscopy techniques like STED and single-molecule localization microscopy, Unclearing Microscopy does not rely on switching fluorescent molecules between different states. Hence, Unclearing Microscopy does not require complex instrumentation or image processing to reveal nanoscopic features in a sample. While this has already been the case for conventional Expansion Microscopy, Unclearing Microscopy completely decouples super-resolution optical microscopy from fluorescence as the means of contrast.

This unprecedented strength enables a wide range of future applications of Unclearing Microscopy. In contrast to fluorescence-based methods, Unclearing Microscopy is not constrained by rate-limited fluorescence photon flux but can draw from essentially unlimited photon rates enabling, in principle, the acquisition of large fields of view (FOV) in less than a millisecond, only limited by available data acquisition rates. Photobleaching, an omnipresent constraint in fluorescence microscopy, is equally negligible allowing for essentially unlimited signal-to-noise ratio imaging of large 3D volumes. We envision Unclearing Microscopy having a large impact in applications requiring examination of tissue ultrastructure in large volumes. For example, combined with instantaneous large-FOV 3D data acquisition offered by next-generation OI-DIC microscopes, PBA-based unclearing may pave the way to moonshot light-based brain connectomics projects since, in principle, it solves the limitations of data acquisition speed and two-dimensionality that EM methods suffer from (Lichtman et al., 2014). Perhaps reflected light confocal microscopy will become prevalent again (Boyde, 1985).

In clinical applications, we see unclearing by chromogenic deposition and direct visualization of the sample potentially replacing microscopic evaluation of H&E-stained clinical samples in the future. More importantly, it offers >10-fold higher resolution imaging of sample ultrastructure when combined with simple compound microscopes, which remain the workhorse histopathology instruments of the healthcare industry (Fine, 2015).

Moreover, for PBA-based unclearing, we also see an opportunity for long-term archiving of patient samples. PBA-uncleared samples represent densely crosslinked polyacrylamide, with very stable properties at room temperature and ambient light conditions (Caulfield et al., 2003).

Finally, our work represents the first recorded instance of seeing cellular microstructures by the naked eye and dispels the centuries-old conviction that magnifying lenses need to stand in between the observer and the observed. Out of the province of science and into the institutions of society and education, Unclearing Microscopy represents an opportunity for the general public to experience material phantoms of the building blocks that make us.

## Methods

Please see **Supplementary Tables 1-2** for an overview of the reagents and materials used in this work.

### Coverslip preparation

Before plating HeLa and COS-7 cells, 12-mm round glass coverslips (Electron Microscopy Sciences, catalog no. 72230-01) were cleaned in a sonic bath (Bronson) submerged in 1 M KOH (Macron Fine Chemicals; catalog no. 6984-04) for 15 min and then rinsed with MilliQ water three times. Coverslips were then sterilized with 100% ethanol and rinsed with sterile phosphate-buffered saline (PBS; Gibco, catalog no. 10010023) before adding media and cells.

### Cell culture

Cells were grown in Dulbecco’s modified Eagle medium (DMEM; Gibco, catalog no. 21063029), supplemented with 10% fetal bovine serum (FBS; Gibco, catalog no. 10438026), and 1% mL/L penicillin-streptomycin (Gibco, catalog no. 15140122) at 37°C with 5% CO2. Cells were passaged twice to three times a week and used between passage number 2 and 20. Passaging was performed using 1× PBS and 0.05% Trypsin-EDTA (Gibco, catalog no. 25300054). Approximately 24 h before fixation, cells were seeded on glass coverslips at ~65,000 cells per well.

### Cell fixation

HeLa cells were fixed with 3% formaldehyde (FA) and 0.1% glutaraldehyde (GA) (Electron Microscopy Sciences, catalog nos. 15710 and 16019, respectively) in 1× PBS for 15 min at RT. COS-7 cells were fixed with 4% FA + 0.1% GA in 1× PBS for 15 min at RT. Samples were rinsed three times with 1× PBS and processed according to the pan-ExM protocol immediately after.

### Expansion reagents

Acrylamide (AAm; catalog no. A9099), N,N’-(1,2-dihydroxyethylene)bisacrylamide (DHEBA; catalog no. 294381), N,N’-Cystaminebisacrylamide (BAC; catalog no. 9809), and sodium hydroxide (NaOH; catalog no. S8045) were purchased from Sigma-Aldrich. Sodium acrylate (SA) was purchased from Santa Cruz Biotechnology (catalog no. sc-236893C). 38% (w/v) SA solutions were made and centrifuged at 4000 rpm for 5 min. The supernatant was transferred to a new bottle and stored at 4 °C until use. N,N’-methylenebis(acrylamide) (BIS; catalog no. J66710) was purchased from Alfa Aesar. Ammonium persulfate (APS) was purchased from both American Bio (catalog no. AB00112) and Sigma-Aldrich (catalog no. A3678). N,N,N’,N’-tetramethylethylenediamine (TEMED; catalog no. AB02020), 1 M Tris solution, pH 8 (catalog no. AB14043), 5 M NaCl solution (catalog no. AB1915), and 20% sodium dodecyl sulfate solution (SDS; catalog no. AB01922) were purchased from American Bio. 10X phosphate buffered saline (10X PBS; catalog no. 70011044) was purchased from Thermofisher.

### pan-ExM gelation chamber

The gelation chamber was constructed using a glass microscope slide (Sigma-Aldrich, catalog no. S8400) and two spacers, each consisting of a stack of two no. 1.5 22 × 22 mm coverslips (Fisher Scientific, catalog no. 12-541B) (cells) or singular no. 1.5 22 × 22 mm coverslips (tissue sections). The spacers were superglued to the microscope slide on both sides of the neuron-adhered coverslip, with this latter coverslip glued in between. A no. 1.5 22 × 22mm coverslip was used as a lid after addition of the gel solution. This geometry yielded an initial gel thickness size of ~170 μm for both cells and tissue.

### First round of expansion for HeLa cells

Cells, previously fixed as described in the **Cell fixation** section, were incubated in post-fix solution (0.7% FA + 1% AAm (w/v) in 1× PBS) for 8-10 h at 37°C. Next, cells were washed twice with 1× PBS for 10 min each on a rocking platform and embedded in the first expansion gel monomer solution (18.75% (w/v) SA + 10% AAm (w/v) + 0.1% (w/v) DHEBA + 0.25% (v/v) TEMED + 0.25% (w/v) APS in 1× PBS). Gelation proceeded first for 10-15 min at room temperature (RT) and then for 1.5 h at 37 °C in a humidified chamber. Coverslips with hydrogels were then incubated in ~4 mL denaturation buffer (200 mM SDS + 200 mM NaCl + 50 mM Tris in MilliQ water, pH 6.8) in 6-well plates for 30 min at 37°C. Gels were then transferred to denaturation buffer-filled 1.5 mL Eppendorf tubes and incubated at 76 °C for 1 h. Next, the gels were washed twice with PBS for 20 min each and optionally stored in PBS overnight at 4 °C. Gels were then cut and placed in 6-well plates filled with MilliQ water for the first expansion. Water was exchanged twice every 30 min and once for 1 h. Gels expanded between 4.5× and 5.0× according to SA purity.

### First round of expansion for brain tissue sections

70 μm-thick mouse brain tissue sections from wild-type (C57BL/6) male mice between P21 and P35 were prepared as described in our earlier work (M’Saad et al., 2022). All experiments were carried out in accordance with National Institutes of Health (NIH) guidelines and approved by the Yale IACUC.

Brain tissue sections were first incubated in inactivated first expansion gel monomer solution (18.75% (w/v) SA + 10% AAm (w/v) + 0.1% (w/v) DHEBA in 1× PBS) for 30-45 min on ice and then in activated first expansion gel monomer solution (18.75% (w/v) SA+ 10% AAm (w/v) + 0.1% (w/v) DHEBA + 0.075% (v/v) TEMED + 0.075% (w/v) APS in 1× PBS) for 15-20 min on ice before placing in gelation chamber. The tissue sections were gelled for 15 min at RT and 2 h at 37°C in a humidified chamber. Next, the tissue-gel hybrids were peeled off of the gelation chamber and incubated in ~4 mL denaturation buffer (200 mM SDS + 200 mM NaCl + 50 mM Tris in MilliQ water, pH 6.8) in 6-well plates for 15 min at 37 °C. Gels were then transferred to denaturation buffer-filled 1.5 mL Eppendorf tubes and incubated at 76 °C for 4 h. The gels were washed twice with 1× PBS for 20 min each and once overnight at RT. Gels were optionally stored in 1× PBS at 4 °C. The samples were then placed in 6-well plates filled with MilliQ water for the first expansion. Water was exchanged three times every 1 h. Gels expanded between ~5.5× and ~6.0× according to SA purity.

### Re-embedding in neutral hydrogel

Expanded hydrogels (of both cell and brain tissue samples) were incubated in fresh re-embedding neutral gel monomer solution (10% (w/v) AAm + 0.05% (w/v) DHEBA + 0.05% (v/v) TEMED + 0.05% (w/v) APS) two times for 20 min each on a rocking platform at RT. Immediately after, the residual gel solution was removed by gentle pressing with Kimwipes. Each gel was then sandwiched between one no. 1.5 coverslip and one glass microscope slide. Gels were incubated for 1.5 h at 37 °C in a nitrogen-filled and humidified chamber. Next, gels were detached from the coverslips and washed two times with 1× PBS for 20 min each on a rocking platform at RT. The samples were stored in 1× PBS at 4 °C. No additional post-fixation of samples after the re-embedding step was performed.

### Second round of expansion

For samples that were uncleared with chromogens, re-embedded hydrogels were incubated in fresh second gel monomer solution (18.75% (w/v) SA + 10% AAm (w/v) + 0.1% (w/v) BIS + 0.05% (v/v) TEMED + 0.05% (w/v) APS in 1× PBS) two times for 15 min each on a rocking platform on ice. For samples that were uncleared with PBA, re-embedded hydrogels were incubated in fresh second gel monomer solution (18.75% (w/v) SA + 10% AAm (w/v) + 0.15% (w/v) BAC + 0.05% (v/v) TEMED + 0.05% (w/v) APS in 1× PBS) two times for 15 min each on a rocking platform on ice. Each gel was then sandwiched between one no. 1.5 coverslip and one glass microscope slide. Gels were incubated for 1.5 h at 37°C in a nitrogen-filled and humidified chamber. Next, to dissolve DHEBA, gels were detached from the coverslips and incubated in 200 mM NaOH for 1 h on a rocking platform at RT. Gels were afterwards washed three to four times with 1× PBS for 30 min each on a rocking platform at RT or until the solution pH reached 7.4. The gels were optionally stored in 1× PBS at 4 °C. Subsequently, gels to be uncleared with chromogens were pan-stained with NHS ester-biotin followed by HRP-conjugated streptavidin (as described in the **Unclearing by chromogen deposition** section) and gels to be uncleared with photopolymers were pan-stained with eosin-5-isothiocyanate (as described in the **Unclearing by photopolymerization** section). Gels were extensively washed with 0.1% TX-100 in PBS (0.1%PBS-T) and placed in 6-well plates filled with MilliQ water for the second expansion. Water was exchanged at least three times every 1 h at RT. Gels expanded to a final expansion factor of ~14× to ~20× according to the crosslinker used (cells) and 23× (brain tissue) and sectioned to a thickness of 1 mm on a Leica VT1000 S vibrating blade microtome.

### Unclearing reagents

N-methyldiethanolamine (MDEA; catalog no. 471828), N-vinyl-2-pyrrolidone (VP; catalog no. V3409), Evans Blue (catalog no. E2129), Direct Red 81 (catalog no. 195251), and eosin-5-isothiocyanate (catalog no. 45245), sodium bicarbonate (catalog no. SLBX3650), and Triton X-100 (TX-100; catalog no. T8787) were purchased from Sigma-Aldrich. NHS-PEG4-Biotin (catalog no. 21330) and HRP-conjugated streptavidin (catalog no. N100) were purchased from Thermofisher. DAB (catalog no. ab64238) and DAB (catalog no. 34002) were purchased from abcam and Thermofisher, respectively. EnzMet (catalog no. #6001-30ML) was purchased from Nanoprobes.

### Unclearing by chromogen deposition

Gels were incubated overnight with 50 μM NHS-PEG4-Biotin dissolved in 1× PBS on a rocking platform at 4 °C. The gels were subsequently washed five times in 0.1%PBS-T for 20 min each on a rocking platform at RT. Gels were incubated with 1 μg/mL HRP-conjugated streptavidin in 1× PBS overnight on a rocking platform at RT, washed five times in 0.1%PBS-T for 20 min each, expanded to its maximum size in MilliQ water, and sectioned with a vibratome as detailed in **Second round of expansion** section. Gels were next uncleared with either DAB or EnzMet until visible (between 3 and 20 min) and according to kit instructions. Unclearing was stopped by two 5-min MilliQ water exchanges.

### Unclearing by photopolymerization

After polymerization-based signal amplification, gels were incubated overnight with 200 μM eosin-5-isothiocyanate dissolved fresh in 100 mM sodium bicarbonate solution on a rocking platform at 4°C. The gels were subsequently washed three to five times in 0.1%PBS-T for 20 min each on a rocking platform at RT and washed once overnight at 4°C.

After eosin-5-isothiocyanate pan-staining, the gels were incubated in fresh monomer solution (38% AAm (w/v) + 2% BIS (w/w) + 210 mM MDEA + 35 mM VP) two times for 10 min each on a rocking platform at RT. The samples were then irradiated with 530 nm collimated LED light (QUADICA, catalog no. SP-03-G4) at 55 mW/cm^2^ for 10-20 min or until a photopolymer was visible (HeLa cells) and for ~1-2 min or until a photopolymer was visible (brain tissue). The samples were optionally incubated in 1 mg/mL Evans Blue or 2 mg/mL Direct Red 81 for 20 min on a rocking platform at RT and subsequently rinsed with MilliQ water.

### NHS ester ATTO532 pan-staining

After unclearing by photopolymerization, gels were incubated for 1.5 h with 20 μg/mL NHS ester-ATTO532 (Sigma-Aldrich, catalog no. 88793) dissolved in 100 mM sodium bicarbonate solution on a rocking platform at RT. The gels were subsequently washed three to five times in PBS-T for 20 min each on a rocking platform at RT.

### Evans Blue hydrogel staining assay

The photopolymer in **Fig. 2b** is composed of 210 mM MDEA + 35 mM VP + 38% (w/v) AAm + 0.1% (w/v) BIS + 12.5 μM eosin-5-isothiocyanate. The photopolymer in **Fig. 2c** is composed of 210 mM MDEA + 35 mM VP + 10% (w/v) AAm + 19% SA (w/v) + 0.1% (w/v) BIS + 12.5 μM eosin-5-isothiocyanate. The photopolymer in **Fig. 2d** is composed of 210 mM MDEA + 35 mM VP +38% (w/v) AAm + 2% (w/w) BIS + 12.5 μM eosin-5-isothiocyanate. The photopolymer in **Fig. 2e** is composed of 210 mM MDEA + 35 mM VP + 10% (w/v) AAm + 19% (w/v) SA + 2% (w/w) BIS + 12.5 μM eosin-5-isothiocyanate. All photopolymers were photopolymerized with 530 nm collimated LED light at 12 mW/cm^2^ for 10 min on 30-mm glass MatTek dishes (MatTek life science, catalog no. P50G-1.5-30-F) and stained with 1 mg/mL Evans Blue for 20 min on a rocking platform at RT. Photopolymers were then extensively rinsed with deionized water and photographed with an iPhone 11 camera.

### Eosin threshold concentration assay

All photopolymers in **Supplementary Fig. 2** are composed of 210 mM MDEA + 35 mM VP + 38% (w/v) AAm + 2% (w/w) BIS + eosin-5-isothiocyanate at the indicated concentrations. All photopolymers were polymerized with 530 nm collimated LED light on 30-mm glass MatTek dishes at irradiation intensities and durations described in the figure legend.

### Expansion factor calculation in HeLa cells

Using 2-pixel thick line profiles in FIJI, the peak-to-peak distances between NHS ester pan-stained mitochondria cristae were measured and divided by 85 nm, its previously determined value (M’Saad and Bewersdorf, 2020). intensity distributions between Homer1 and PSD, PSD-95 and DP, and Bassoon and PSD were measured and divided by the determined experiment expansion factor. For cells uncleared by chromogenic deposition shown in **Fig. 1**, 82 line profiles were drawn in 25 mitochondria in 5 cells. For cells uncleared by PBA in **Fig. 2i–m**, 83 line profiles were drawn in 35 mitochondria in 7 cells. For cells uncleared by PBA in **Fig. 2n–s**, 90 line profiles were drawn in 30 mitochondria in 7 cells. Expansion factors are reported in the figure legends.

### Expansion factor calculation in brain tissue

Using 2-pixel thick line profiles in FIJI, the peak-to-peak distances between the intensity distributions of the DP and PSD NHS ester signals were measured and divided by 81.9 nm, the DP-PSD length determined in an earlier report (M’Saad et al., 2022). The expansion value determined for this measurement was used to calculate the DP-PSD length of PBA uncleared synapses in the same experiment. For NHS ester PSD-DP measurements, 91 line profiles were drawn from two independent samples. For PBA uncleared PSD-DP measurements, 63 line line profiles were drawn from two independent samples in the same experiment. Results are shown in **Fig. 3r**.

### Image acquisition

Images shown in **Fig. 1a**, **Fig. 1c–m**, **Fig. 2b–e**, **Fig. 2i–j**, and **Supplementary Figs. 1-4** were acquired using an Apple iPhone 11 camera.

Phase contrast and confocal images were acquired using a Leica SP8 STED 3X equipped with a SuperK Extreme EXW-12 (NKT Photonics) pulsed white light laser as an excitation source. All images were acquired either using a HC PL APO 63×/1.2 water objective. ATTO532 was imaged with 532-nm excitation. Application Suite X software (LAS X; Leica Microsystems) was used to control imaging parameters.

### OI-DIC imaging

OI-DIC (Orientation-Independent Differential Interference Contrast) images were acquired using a custom-built OI-DIC module (Malamy and Shribak, 2018), which is installed on an Olympus BX61 microscope equipped with a 60x/1.2NA water immersion objective lens (UPLSAPO60XW), condenser (U-TLD), and CCD camera (Infinity 3-1M; Lumenera). For illumination, we used an Xcite 200DC light source (Excelitas) and a bandpass filter BP546/10 (Chroma) with a central wavelength of 546 nm and 10-nm bandwidth. The OI-DIC module includes two crossed DIC prisms (U-DICTHR; Olympus) that sheared two interfering beams by 170 nm distance.

To acquire OI-DIC images we capture a set of six raw DIC images with two orthogonal shear directions and three different biases. The obtained images are used to compute a quantitative image (map) of the optical path difference (OPD) gradient vector, which is then integrated in order to create an OPD map. The OPD equals the product of thickness and refractive index difference and represents the cell dry mass. For a more visually informative (yet more qualitative) rendering, we chose to process our OI-DIC data with the inverse Riesz transform. This simple method combines the two orthogonal differential images in a way that maintains the strong edge emphasis, yet enforces isotropy (i.e. rotation invariance). It is equivalent to the sum of a pair of 2-D linear filters separately operating on the two differential images (Larkin & Fletcher, 2014; Shribak *et al*., 2017). The inverse Riesz transform image can be considered to be an OPD image that has edges and small details enhanced. The enhancement is equivalent to a high-pass image processing filter

### Image processing

Images were visualized, smoothed and contrast-adjusted using FIJI/ImageJ software version 2.1.0/.53c. Minimum and maximum brightness were adjusted linearly for optimal contrast. All line profiles were extracted from the images using the Plot Profile tool in FIJI/ImageJ.

### Statistics and reproducibility

For all quantitative experiments, the number of samples and independent reproductions are listed in the figure legends.

## Data availability

The datasets generated and/or analyzed during the current study are available from the corresponding author on reasonable request.

## Acknowledgements

We would like to acknowledge Dr. Xinran Liu for the TEM image. We would like to thank Phylicia Kidd for assistance with cell culture, Dr. Ilona Kondratiuk and Dr. Yifei Cai for providing mouse brain sample sections, and Hannahmariam Mekbib for LED light collimation. We would like to thank Dr. Kenny Chung, Dr. James Rothman, Dr. Derek Toomre, and Dr. Zach Marin for important discussions. This work was supported by grants from the Wellcome Trust (203285/B/16/Z) and NIH (P30 DK045735, S10 OD020142). M.S. gratefully acknowledged funding from the NIGMS/NIH (R01 GM101701) and from the Inoue Endowment Fund.

## Author contributions

O.M. developed sample preparation protocols. O.M. and M.S. performed OI-DIC imaging. O.M. and J.B. interpreted the data. O.M. and J.B. wrote the manuscript.

