## Supplementary Tables and Figures for "Unclearing Microscopy"

### SUPPL. TABLE 1

**Supplementary Table 1:** Reagents used in fixation, polymerizations, and unclearing

| Step | Reagent | Acronym | Storage | Catalog number | Vendor |
|---|---|---|---|---|---|
| Polymerizations | 40% Acrylamide | AAm | 4°C | A9099 | Sigma-Aldrich |
| | N,N'-(1,2-Dihydroxyethylene)bisacrylamide | DHEBA | 4°C | 294381 | Sigma-Aldrich |
| | N,N′-Cystaminebisacrylamide | BAC | -20°C desiccated | 9809 | Sigma-Aldrich |
| | Sodium acrylate | SA | -20°C desiccated | sc-236893C | SCBT* |
| | N,N′-Methylenebis(acrylamide) | BIS | RT | J66710 | Alfa Aesar |
| | Ammonium persulfate | APS | RT desiccated | AB00112 | American Bio |
| | Ammonium persulfate | APS | RT desiccated | A3678 | Sigma-Aldrich |
| | N,N,N′,N′-Tetramethylethylenediamine | TEMED | RT desiccated | AB02020 | American Bio |
| Unclearing | N-methyldiethanolamine | MDEA | RT | 471828 | Sigma-Aldrich |
| | N-vinyl-2-pyrrolidone | VP | RT | V3409 | Sigma-Aldrich |
| | Evans Blue | EB | RT | E2129 | Sigma-Aldrich |
| | Direct Red 81 | - | RT | 195251 | Sigma-Aldrich |
| | Eosin-5-isothiocyanate | Eosin | 4°C | 45245 | Sigma-Aldrich |
| | NHS-PEG4-Biotin | - | -20°C desiccated | 21330 | Thermofisher |
| | HRP-conjugated streptavidin | HRP-ST | 4°C | N100 | Thermofisher |
| | DAB Substrate Kit | - | 4°C | 34002 | Thermofisher |
| | DAB Substrate Kit | - | 4°C | ab64238 | abcam |
| | EnzMet™ IHC / ISH HRP Detection Kit | - | 4°C | 6001-30ML | Nanoprobes |
| Buffers | Sodium hydroxide | NaOH | RT | S8045 | Sigma-Aldrich |
| | 20% Sodium dodecyl sulfate solution | SDS | RT | AB01922 | American Bio |
| | 5 M NaCl solution | NaCl | RT | AB1915 | American Bio |
| | 1 M Tris solution, pH 8 | Tris | RT | AB14043 | American Bio |
| | Triton X-100 | TX-100 | RT | T8787 | Sigma-Aldrich |
| | 1X Phosphate buffered saline (Gibco) | 1X PBS | RT | 10010023 | Thermofisher |
| | 10X Phosphate buffered saline (Gibco) | 10X PBS | RT | 70011044 | Thermofisher |
| | Sodium Bicarbonate | - | RT | S5761 | Sigma-Aldrich |
| Fixatives | 16% Paraformaldehyde | FA | RT | 15710 | EMS** |
| | 8% Glutaraldehyde | GA | 4°C | 16019 | EMS** |

* Santa Cruz Biotechnology; ** Electron Microscopy Sciences

### SUPPL. TABLE 2

**Supplementary Table 2:** Materials used

| Materials | Vendor | Catalog number |
|---|---|---|
| No. 1.5 12-mm round glass coverslips | Electron Microscopy Sciences | 72230-01 |
| Glass microscope slide | Sigma-Aldrich | S8400 |
| No. 1.5 22 x 22 mm square cover glass coverslips | Fisher Scientific | 12-541BP |
| No. 1.5 18-mm round coverslip | Marienfeld | 0117580 |
| 50 mm MatTek dish, No. 1.5 coverslip 30-mm diameter | MatTek Life Sciences | P50G-1.5-30-F |
| Green (530 nm) LED | QUADICA | SP-03-G4 |

SUPPL. FIG. 1

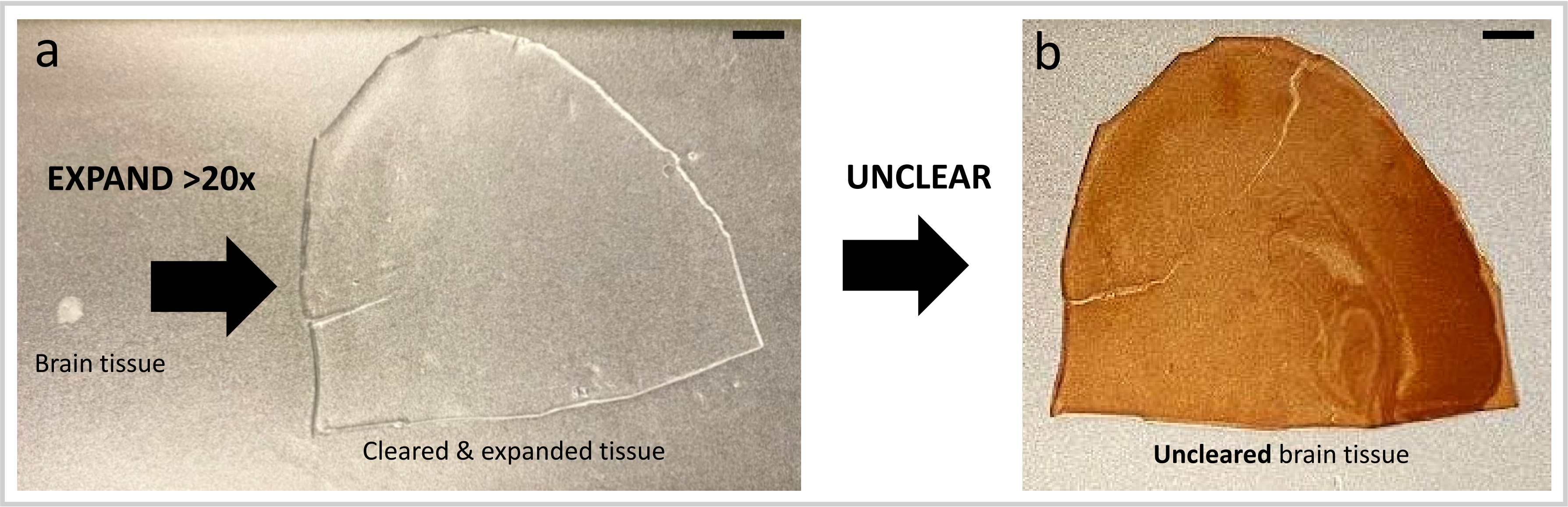


**Supplementary Figure 1: Process of HRP based unclearing of a mouse brain tissue section. a**, Mouse brain tissue section before and after ~24-fold linear sample expansion. **b**, Mouse brain tissue section after HRP-DAB based unclearing. Images were acquired on a cell phone camera. Scale bars are not corrected for the expansion factor. Scale bars (**a**, **b**) 1 cm.

### SUPPL. FIG. 2

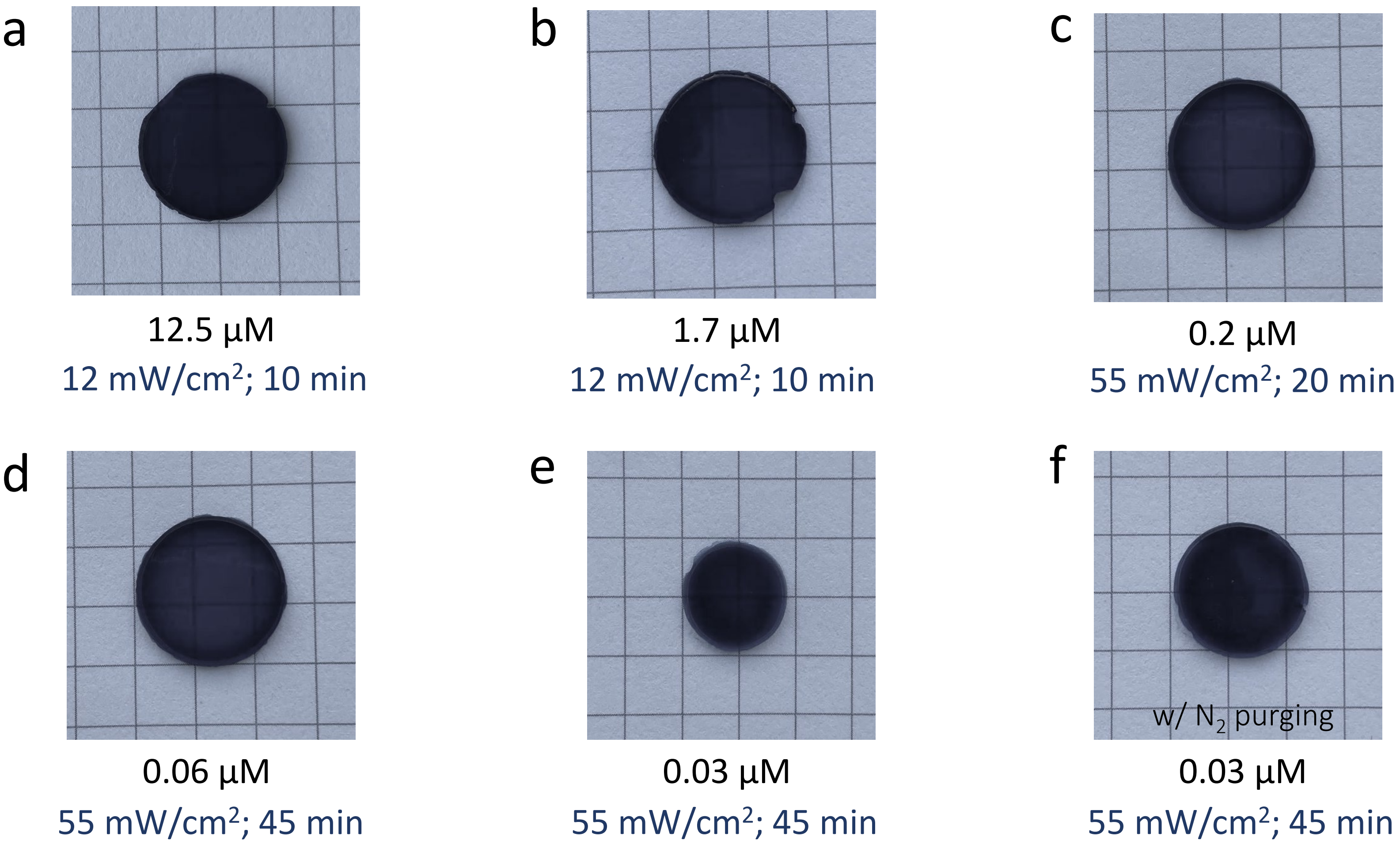


g

| **Distance between two proteins** | **2D density** | **2D density (post-20x pan-ExM)** | **Molar concentration** | **Molar concentration (post-20x pan-ExM)** |
|---|---|---|---|---|
| 5 nm | 40,000 per µm$^2$ | 100 per µm$^2$ | 100 mM | 12.5 µM |
| 10 nm | 10,000 per µm$^2$ | 25 per µm$^2$ | 13 mM | 1.7 µM |
| 20 nm | 2,500 per µm$^2$ | 6.25 per µm$^2$ | 1.7 mM | 0.2 µM |
| 30 nm | 1,111 per µm$^2$ | 2.78 per µm$^2$ | 0.5 mM | 0.06 µM |
| 40 nm | 625 per µm$^2$ | 1.56 per µm$^2$ | 0.2 mM | 0.025 µM |

Threshold eosin surface density = **2.8 eosins per µm$^2$** (Avens et al. 2008)

**Supplementary Figure 2: Eosin threshold concentration assay. a-f**, Photopolymers composed of 38% (w/v) AAm + 2% (w/w) BIS + 210 mM MDEA + 35 mM VP + [indicated concentration] eosin-5-isothiocyanate, photopolymerized with 530 nm LED light with [indicated intensity and exposure time] and stained with 1 mg/mL Evans Blue dye in PBS. **g**, Table summarizing the 2D densities and molar concentrations of proteins in a typical HeLa cell pre- and post-expansion had they been separated 5, 10, 20, 30 or 40 nanometers from one another.

### SUPPL. FIG. 3

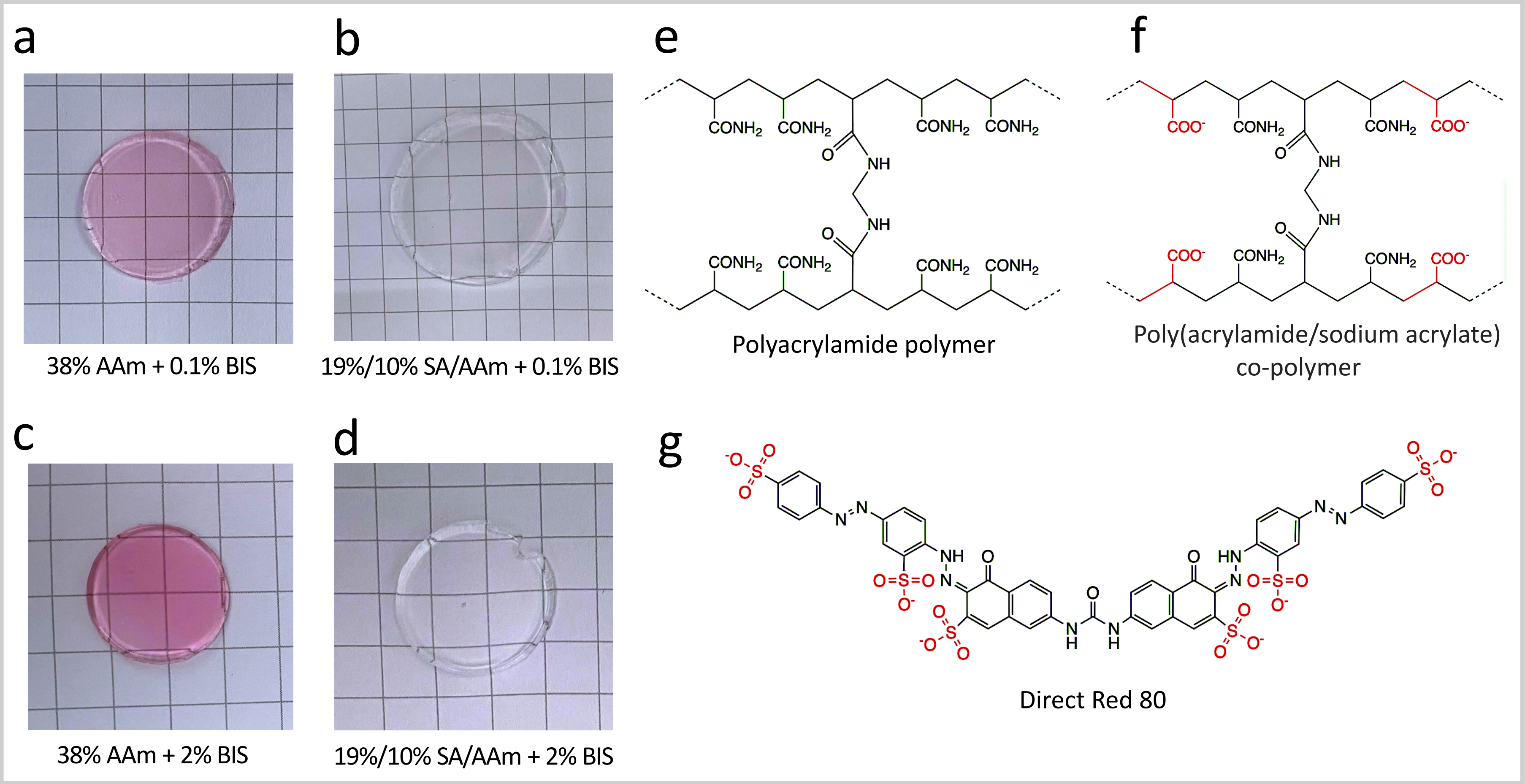


**Supplementary Figure 3: Justification of using Direct Red 81 in staining PBA uncleared samples. a-d,** Direct Red 81, an anionic dye, efficiently stains neutral polyacrylamide hydrogels (**a** and **c**) but not anionic poly(acrylamide/sodium acrylate) co-polymers (**b** and **d**), regardless of hydrogel crosslinker concentration. **e**, Chemical structure of polyacrylamide (pAAm) polymer. **f**, Chemical structure of poly(acrylamide/sodium acrylate) co-polymer (pAAm/SA) (red: negatively charged carboxylic groups). **g**, Chemical structure of Direct Red 81 dye (red: negatively charged sulfate groups that repel carboxylic groups in anionic pAAm/SA hydrogels and prevent staining of the hydrogel).

SUPPL. FIG. 4

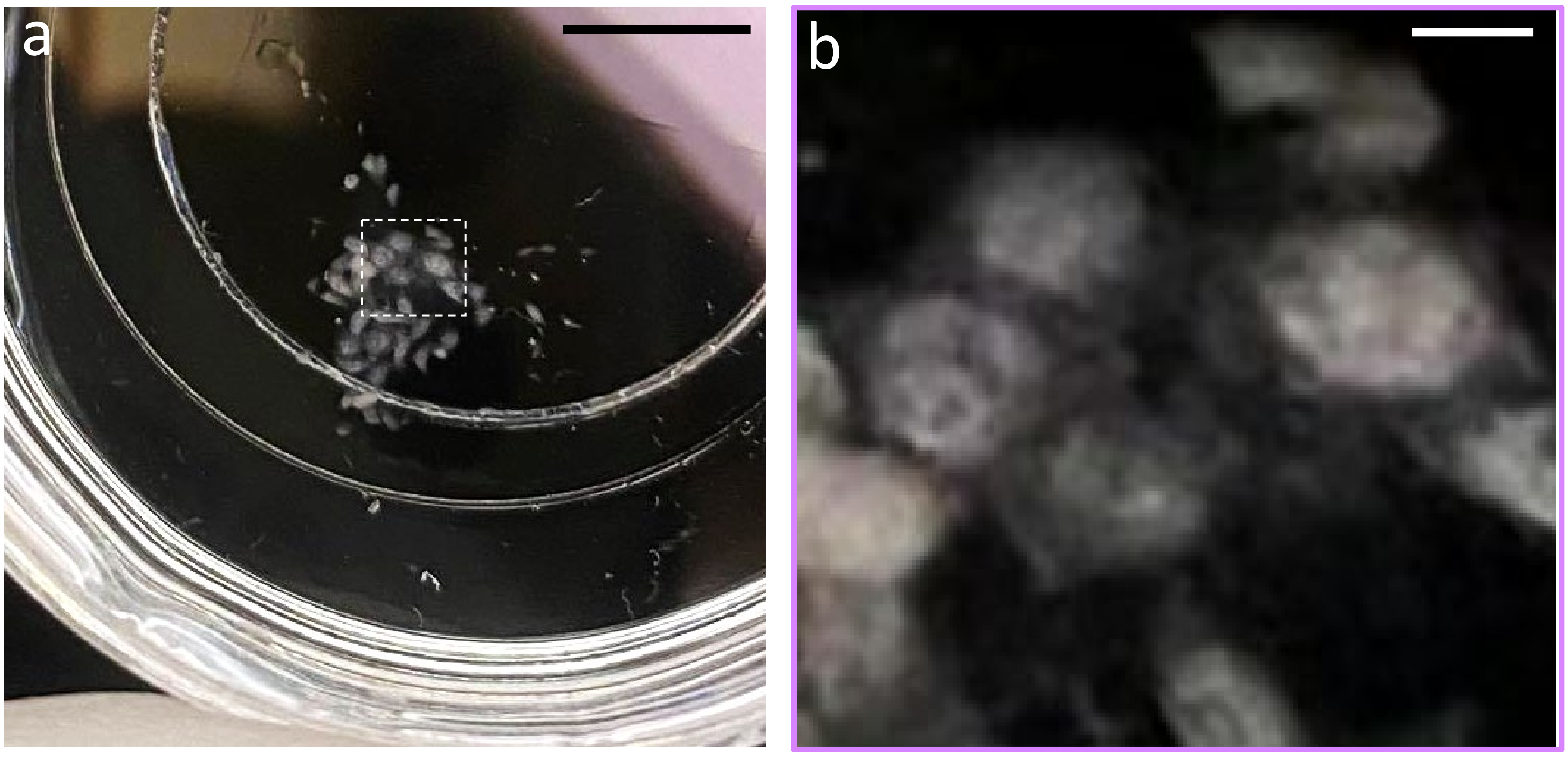


**Supplementary Figure 4: PBA uncleared and non-stained HeLa cells are visible behind a black background.** **a**, PBA uncleared and unstained HeLa cells imaged with a cell phone camera. **b**, Magnified view of the area outlined by the dashed white box in **a**. Scale bars are not corrected for the expansion factor. Scale bars, (**a**) 5 mm, (**b**) 0.5 mm

SUPPL. FIG. 5

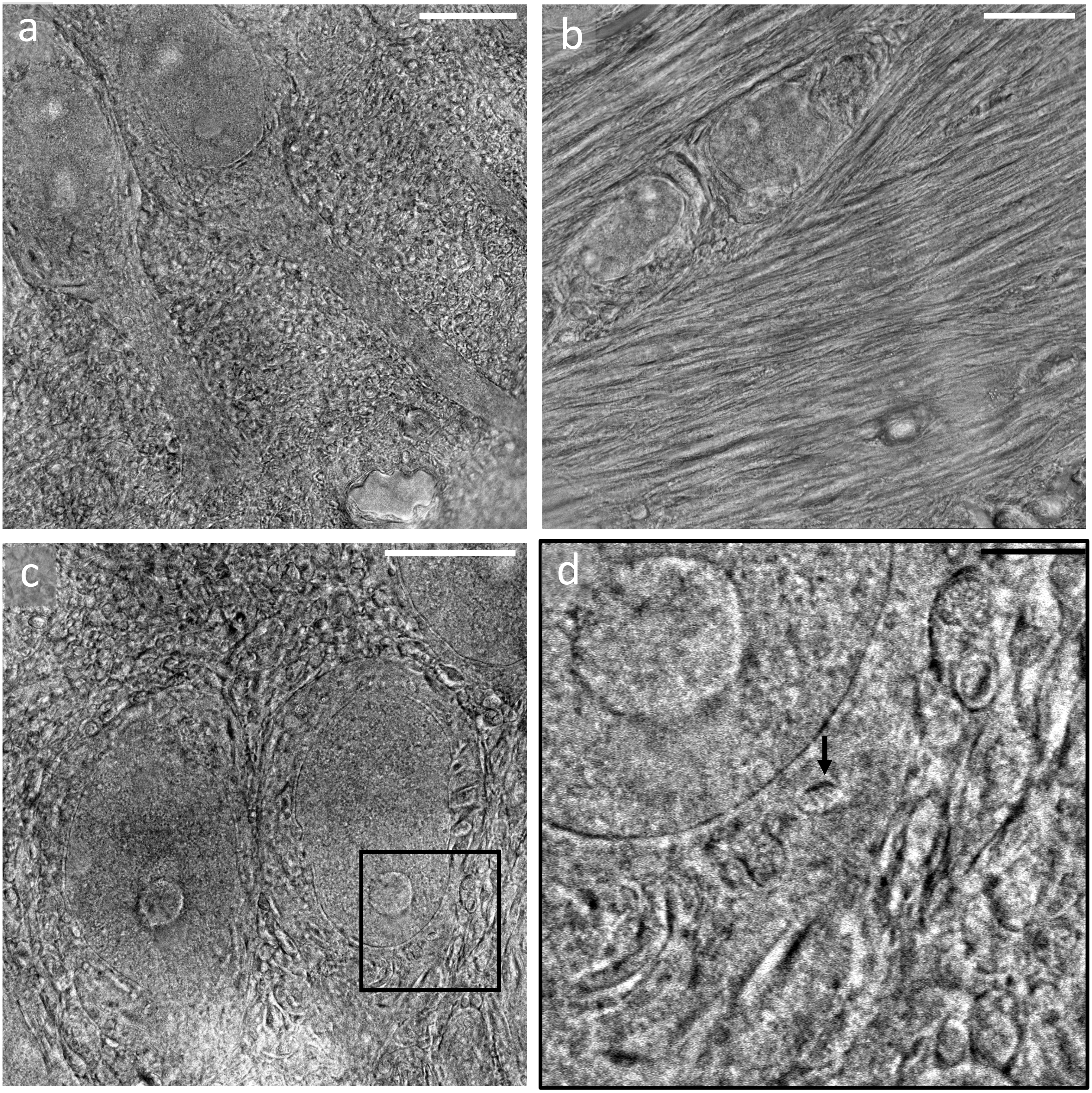


**Supplementary Figure 5: PBA uncleared mouse brain tissue samples imaged with conventional phase contrast microscopy.** **a**, **c**, PBA uncleared cortical neurons. **b**, PBA uncleared axons. **d**, Magnified view of the area outlined by the black box in **c** showing ultrastructural features like mitochondria cristae (arrow). Thin neurites and synapses cannot be distinguished because of poor optical sectioning. Scale bars are not corrected for the expansion factor. Scale bars (**a**-**c**) 5 µm, (**d**) 1 µm.
